# A dynamic humidity arena to explore humidity related behaviours in insects

**DOI:** 10.1101/2023.10.20.563253

**Authors:** Ganesh Giri, Nicolas Nagloo, Anders Enjin

## Abstract

Humidity is a critical environmental factor influencing the behaviour of terrestrial organisms. Despite its significance, the neural mechanisms and behavioural algorithms governing humidity sensation in insects remain elusive. In this study, we introduce a novel dynamic humidity arena to investigate humidity-guided behaviour in the vinegar fly *Drosophila melanogaster*. The arena allows precise humidity control, low error rates, and fast settling times, making it a robust tool for studying humidity-related behaviours. Our results reveal that desiccated and starved flies (DS flies) search for higher relative humidity environments (65-75%) while sated flies tend to stay within their initial environments. In contrast, Ionotropic receptor (Ir)93a mutant flies with impaired humidity sensing show no preference to relative humidity. The search for higher humidity in DS flies is reflected in their relatively high displacement and walking speed compared to control and mutant flies. Our novel method manipulates humidity cues to create complex humidity landscapes that respond in real-time to insect movement. This will help us shed light on how humidity shapes behaviour and offers a foundation for further research in the field of hygrosensation.

## Introduction

Humidity is an environmental factor that plays a fundamental role in shaping the ecology and behaviour of land-living organisms^1,2^. Particularly in small poikilothermic animals such as insects are sensitive to variations in humidity and need to navigate to optimal microclimates for survival^3^. Insects also use humidity as cues to guide them in specific behaviours such as host-seeking and oviposition site-selection ^4–6^. Despite the vital roles of humidity sensation, the neural mechanisms and behavioural algorithms underlying this sense remain unclear.

Insects perceive humidity via specific humidity receptor neurons (HRNs). These HRNs are situated within the hygrosensilla, a structure located inside the antennae on their head^7^. In the case of the vinegar fly *Drosophila melanogaster*, these HRNs are located in an invagination on the posterior side of the antenna, referred to as the sacculus^8-12^. Within the hygrosensilla there are three distinct types of neurons working in unison, forming a hygrosensory triad. These neurons include one moist neuron (responding to increase in humidity), one dry neuron (responding to decrease in humidity and one hygrocool neuron (responding to cooling but not required for temperature preference behaviour). These neurons depend on a combination of ionotropic receptors (IRs) for their function, with IR25a and IR93a expressed and required in all HRNs and IR40a and IR68a specific for subtypes of HRNs^7–10^.

Previous experiments in *Drosophila*, have shown that within the genus, the preferred relative humidity (RH) of species can vary by wide margins. *D. mojavensis*, which inhabits the arid Sonoran Desert in the southwestern United States and northern Mexico, exhibits a strong preference for exceedingly low humidity levels of around 20% RH^9,13^. In contrast, the rainforest–dwelling species, *D. teissieri*, native to the warm and humid tropical rainforests of western Africa, displays a preference for higher humidity, typically favouring an environment with *around 85% RH*^*9*.^ The vinegar fly *Drosophila melanogaster* prefers a humidity around 70% RH^7,9,11,12^. These great differences in preferred RH seem to coincide with the surrounding climate of each species and suggest that species can adapt their RH preference to their surroundings. In addition to these long-term adaptations, humidity seeking behaviour can be affected on shorter time scales by the internal state of the animals with reports that sated *D. melanogaster* avoid high humidities while desiccated and starved flies prefer a higher humidity ^7,13–15^.

While these studies provide some measures of preferred RH, they are all based on binary choices of fixed humidities that can at most only provide a snapshot of the complex humidity-based behaviours. An assay with high temporal resolution where RH can be manipulated in real time is necessary to determine how insects use humidity sensing to make decisions and how different contexts lead to different humidity-based behaviour. Such kind of setup would also allow us to investigate the neural circuitry which underlies such behaviours. Here, we describe an adapted fly-on-ball assay^16,17^ where flies can freely choose humidity ranging from 10% to 80% RH. Using this assay, we show that sated wild type *w*^*1118*^ and sated Ir93a mutant flies have a low activity level and no significant preference towards humidity, whereas desiccated and starved *w*^*1118*^ flies have a higher activity and prefer a humidity within the range of 65-70%. Desiccated and starved IR93a mutant flies with deficient humidity-sensing have similar activity profile as desiccated and starved *w*^*1118*^ flies but are unable to remain at a certain level of humidity. These results are aligned with previous work done in *D. melanogaster* which validate the operation of our dynamic humidity arena and also provide more details on the effect of humidity on fly behaviour.

## Results

### Characterizing the Technical Performance of the Experimental Setup

To assess humidity-guided behaviour of flies we developed a fly-on-ball assay with dynamic changes in RH controlled by the movement of the fly. The required RH level was generated by mixing dry and moist air stream using proportional valves (Figure 1A). Using this system, we formulated two experimental paradigms; one that allows the tested fly to sample all humidities and one where the fly is forced to a certain humidity.

**Figure 1:**
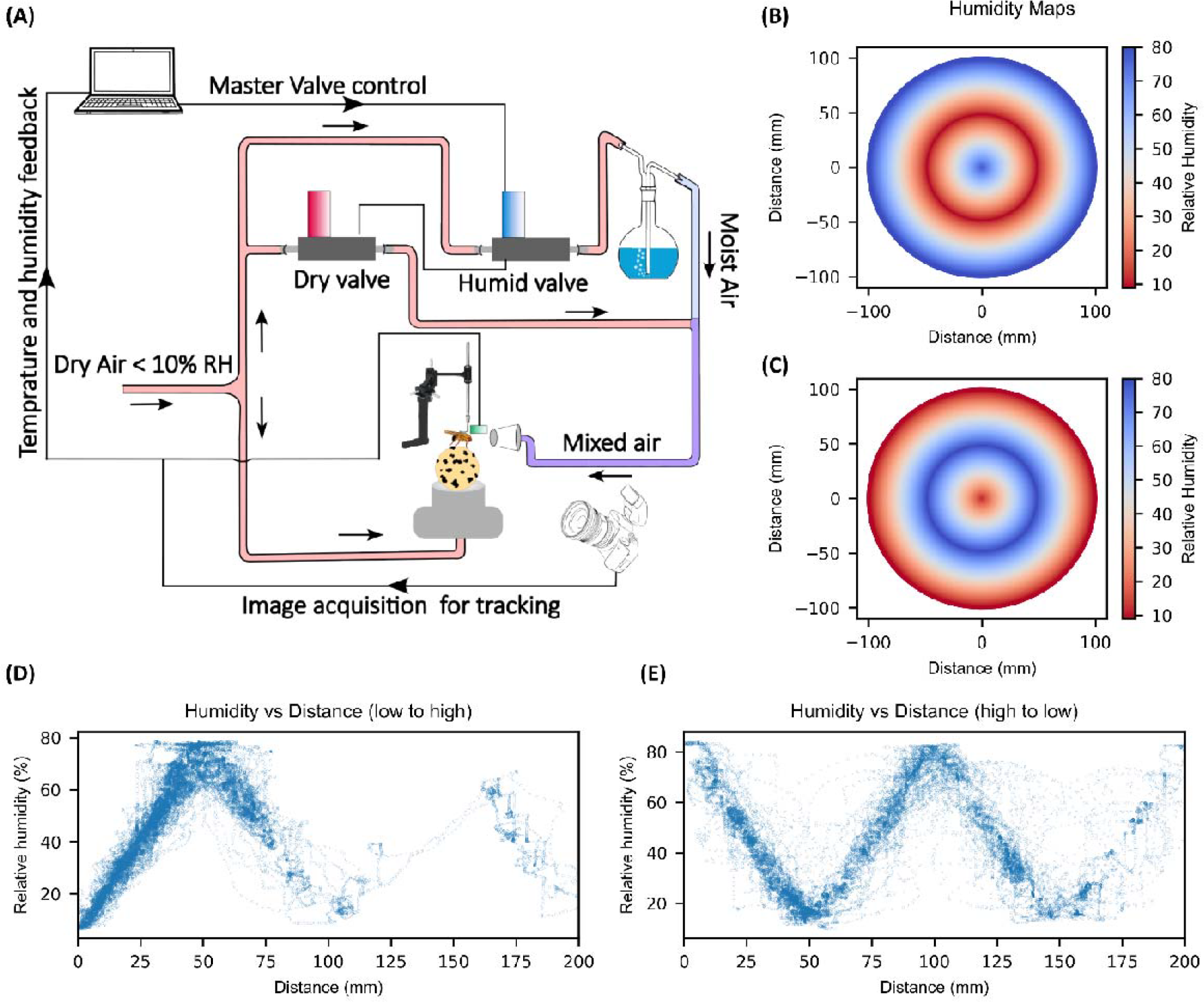
System design and response. Schematic for the dynamic humidity arena (A). The dry and moist air are mixed in required proportions by regulating their flow using the dry and humid valve. to make it a closed loop system, a humidity and temperature sensor placed close to the fly’s antenna sends the feedback response to the computer which makes further adjustments in the valves to minimize the error. A high-speed USB camera is used to acquire the rotation of the ball to calculate the trajectory of the tethered fly. Using the calculated trajectory, humidity values are adjusted depending upon the experimental paradigm. humidity maps where humidity is dependent on the radial distance from the centre. High to low humidity map (B) and low to high humidity map (C). Humidity of tethered fly plotted against the distance travelled from the centre for low to high and high to low humidity map (D-E).

In the first experimental paradigm, we devised a humidity map based on a triangular wave pattern. This pattern exhibited either a radial escalation of RH succeeded by a reduction (Figure 1B), or a radial reduction succeeded by an escalation (Figure 1C) while maintaining a constant slope. Flies moving in this environment demonstrate strong exploratory behaviours and gain access to the full range of humidity values from 10% to 80% depending on the distance travelled from the initial position (Figure 1D, 1E). This allows the tested flies to sample the full range of continuous relative humidities.

In a second paradigm, RH was independent of the movement of the fly but instead was changed every 600 seconds in a stepwise manner across four pre-set RH values: 10%, 30%, 70% and 80% (Figure 2A). Once RH has reached the desired value, the mean error remained close to zero with a root mean squared error of 3.79 (Figure 2B). Latency of humidity changes is relatively low given the nature of the stimulus with 84.65 s for stepping up RH by 20% - 40% across the range while stepping down RH by 20% - 40% has a shorter latency of 82.49 s (Figure 2C, 2D).

**Figure 2:**
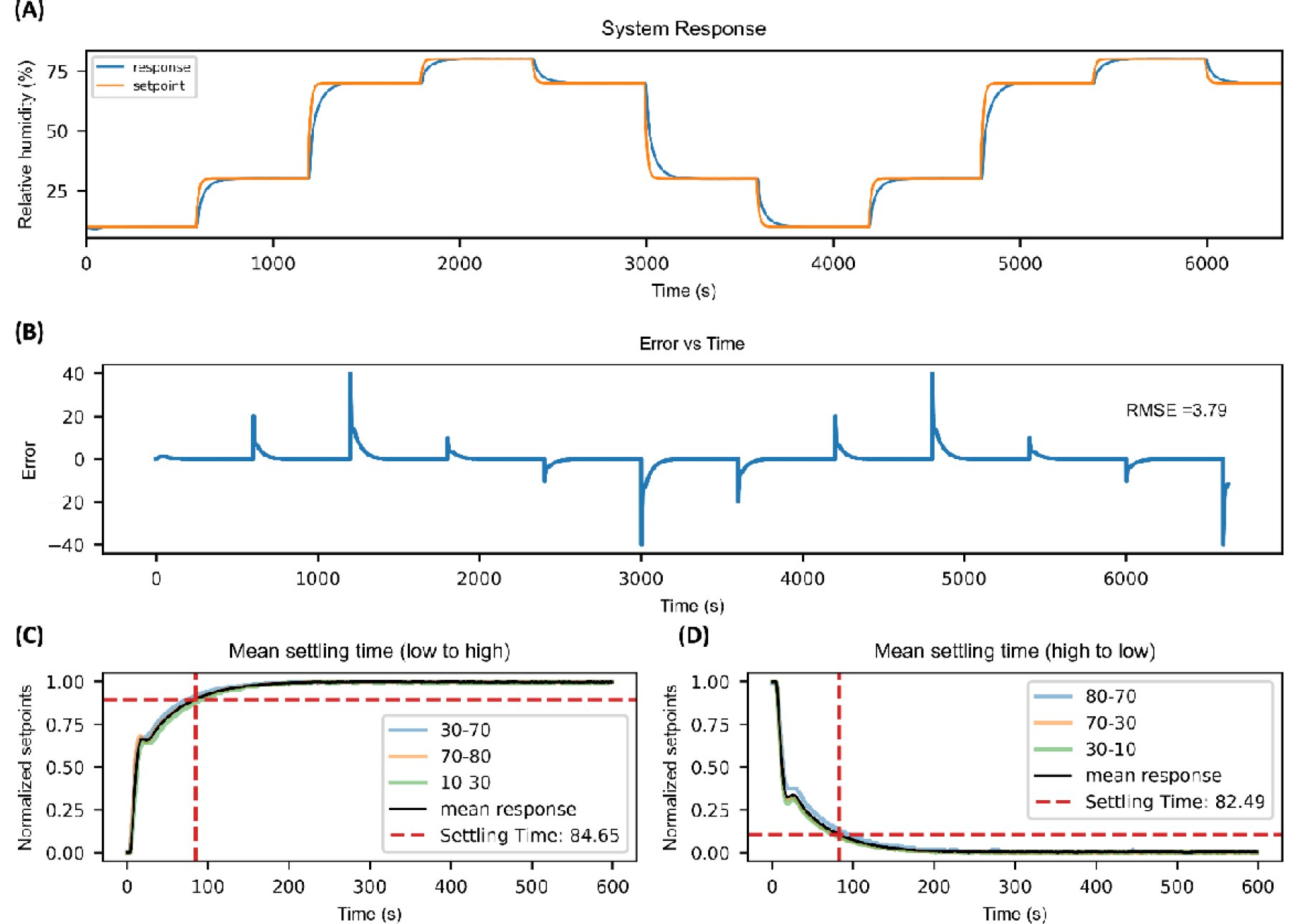
Response of the system to different set points. (A). Error over time between the setpoints and response (B). Settling time for different setpoints. The intercept of the dashed red line along the y axis shows the 90% value of the set point and the intercept along the x axis shows the time taken to achieve the 90% value of the setpoint(C-D).

### Preference Towards Relative Humidity in *D. melanogaster*

To evaluate the preference towards RH of *D. melanogaster*, we used four different groups of flies: desiccated and starved *w*^*1118*^ (DS group) (n=9), sated *w*^*1118*^ (sated group) (n=12), desiccated and starved Ir93a mutants (*Ir93a*^*MI05555*^) deficient in humidity sensing^9,11^ (DS Ir93a group) (n=10) and sated Ir93a mutants (Sated Ir93a group) (n=5). Individuals from each group were allowed to walk on the ball for a duration of one hour. During this period, the RH was automatically adjusted depending on the trajectory of the fly with respect to a pre-set humidity map, either a low-to-high humidity map or a high-to low humidity map. The DS flies exhibited a consistent preference for RH levels ranging from 65% to 70%, irrespective of the humidity map employed (Figure 3A). Conversely, within the sated group, the flies displayed a bimodal preference for RH, dependent on the specific type of humidity map used (Figure 3B). Flies exposed to a low-to-high humidity map favoured RH levels within the range of 15% to 25%, while those exposed to a high-to-low humidity map exhibited a preference for RH levels between 75% and 80%. In contrast, the DS Ir93a group displayed a random distribution of RH preferences (Figure 3C). Sated Ir93a also showed a bimodal humidity preference similar to the sated group depending upon the humidity map employed (Figure 3D). Significant statistical differences were observed in the distribution of humidity preferences among individual flies between the DS group and the rest of the groups (p<0.05) (Figure 3E).

**Figure 3:**
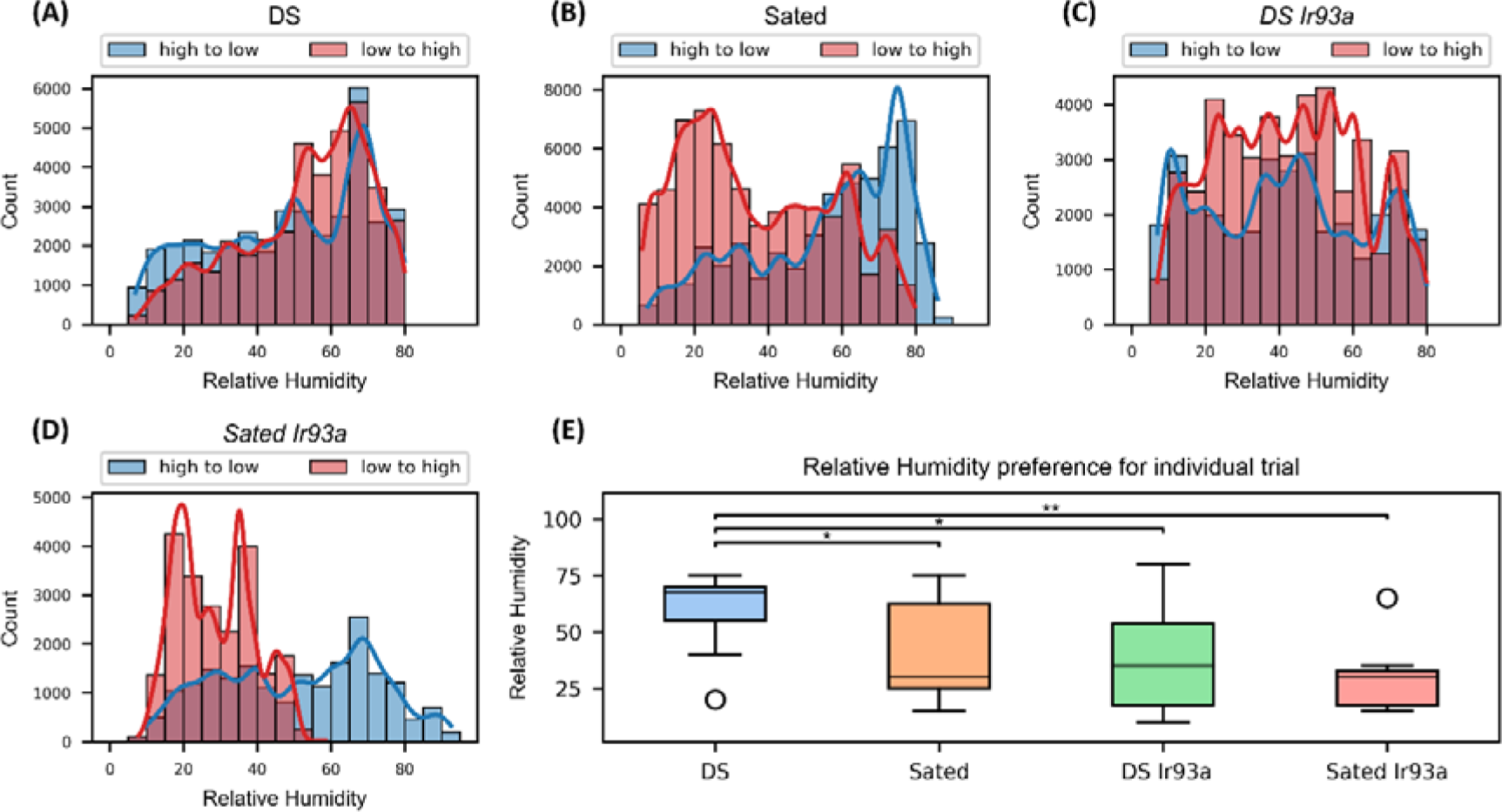
Relative humidity preference. Collective humidity distribution for trials within the DS group, sated group, DS Ir93a and sated Ir93a group (A-D). The peaks in the histogram plot represents the humidity range in which the flies spent the maximum time during observation period. The two different shades used in the histogram represents the type of humidity map used during the experiment. Preferred humidity for individual trial showing statistical significance between the DS group and the remaining groups (D). (** = p<0.01, * = p<0.05)

### Activity of flies in the dynamic humidity arena

Investigating fly activity in terms of displacement and distance covered within the dynamic humidity arena is essential, as it is closely linked to the fluctuations in humidity experienced by the flies. On average, DS flies explored the arena the furthest, covering an average displacement of 205 mm from the arena’s centre (Figure 4A). DS Ir93a flies also displayed substantial exploration, with an average displacement of 142 mm from the centre (Figure 4C). In contrast, sated and sated Ir93a flies exhibited the lowest level of exploration, with an average displacement of 58 mm and 47 mm respectively (Figure 4B and 4D). The total distance covered by the flies followed a similar trend, with both DS and DS Ir93a groups covering significantly greater distances compared to the sated and sated Ir93a group (Figure 4E). Specifically, the median distance covered by DS and DS Ir93a flies was approximately 41% higher than that of the sated and sated Ir93a flies and showed a statistical significance with a p-value of less than 0.05. No statistically significant difference was observed in the total distance covered between the DS and DS Ir93a groups (Figure 4F). RH had an impact on the speed of DS flies. They showed a difference of 0.75 mm/s between its maximum (1.7 mm/s at 20-30% RH) and minimum speed (0.98 mm/s at 60% RH and above). For sated, sated Ir93a *and DS* Ir93a flies the speed remained relatively constant at 0.9 mm/s, 0.6 mm/s and 0.4 mm/s respectively, with no significant variations depending on humidity level (Figure 4G). The predicted speed value using linear mixture model for different RH conditions with flies differentiated based on sex and group showed that there was no difference in speed due to humidity in sated, sated Ir93a and DS Ir93a, however DS flies showed a decline in speed with increasing humidity. Male flies were faster when compared to female flies of the same group (Figure 5A). The assumption of homoscedasticity and normality for the fitted data was met (Figure 5B – 5D).

**Figure 4:**
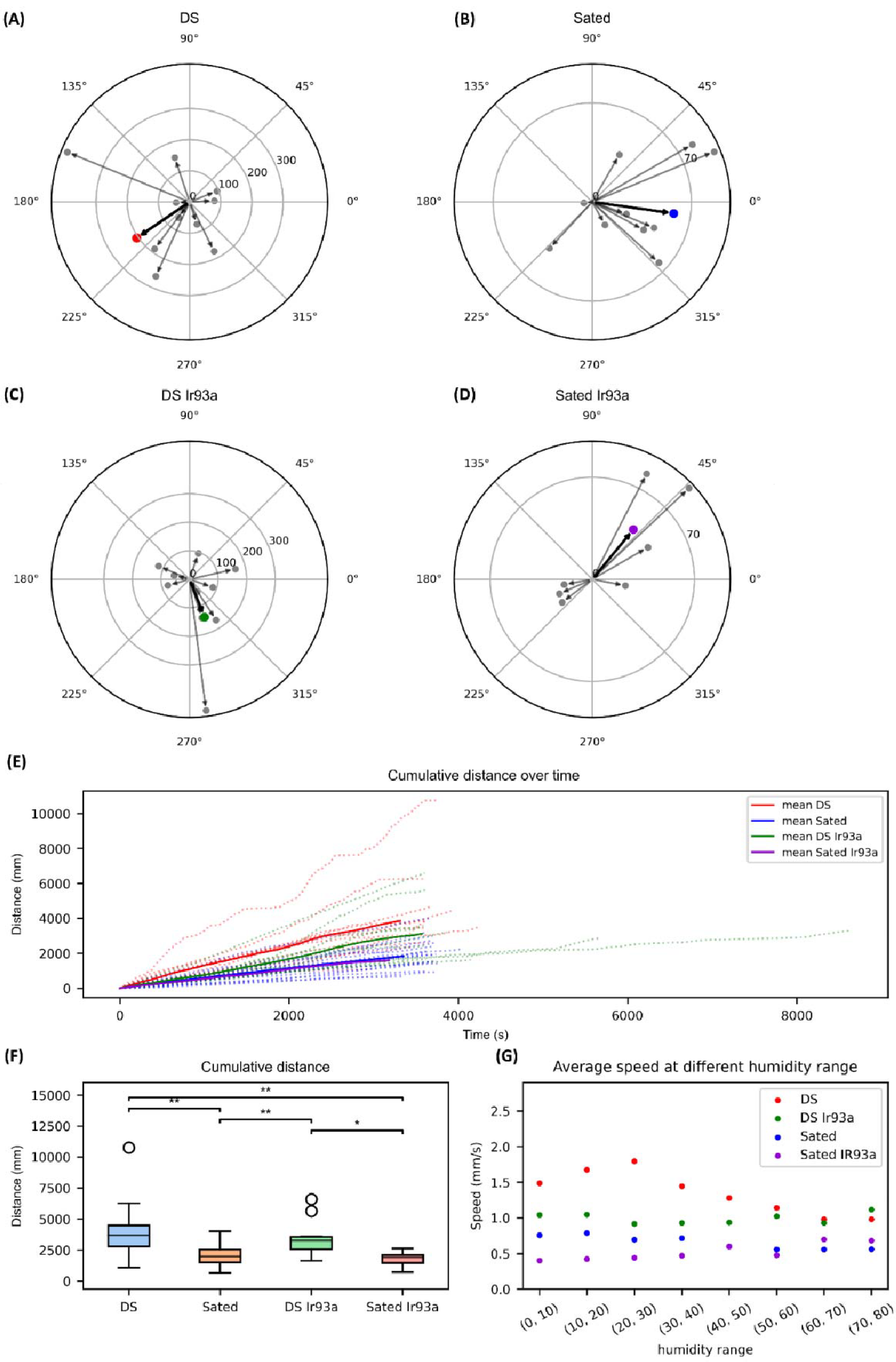
Walking activity in dynamic humidity arena. Maximum displacement of the DS, sated and DS Ir93a and sated Ir93a group represented in the form of polar plot (A-D). The maximum displacement and direction travelled by each fly is represented with an arrow from the centre. The length of the arrow shows the magnitude of displacement. The bold line indicates the mean of maximum displacement within each group. Cumulative distance over time graph showing how the distance travelled changes over time (E). The dashed lines represent individual flies, and the solid line represents the mean of all individuals within the group. Distribution of total distance travelled by each fly within the respective group (F). Average speed of flies grouped at different humidity range (G). (** = p<0.01, * = p<0.05)

**Figure 5:**
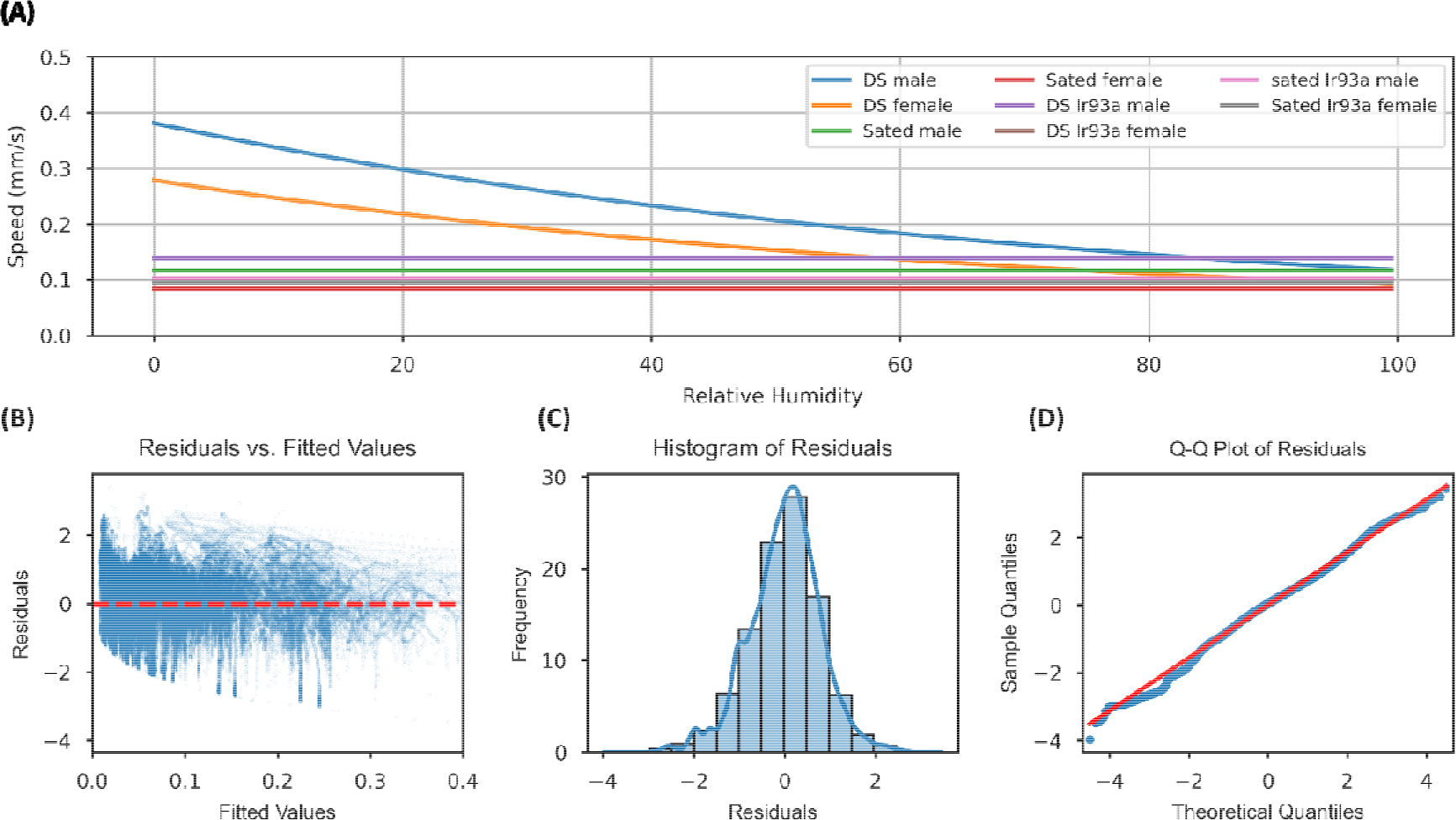
Linear mixed model. Predicted speed for different humidity values of flies, calculated based on group and sex (A). The performance of the model for the provided data is evaluated using the residual vs fitted values (B), the histogram of residuals (D) shows a normal distribution. The QQ plot between the sample quantiles and theoretical quantiles is a straight line showing that they follow a normal distribution (D)

### Experimental validation of speed-humidity relationship

Utilizing the dynamic humidity arena’s capability to control humidity levels for predefined durations, we exposed the four distinct groups to a stepwise humidity alteration, oscillating between 10% and 80% RH, each phase was sustained for 500 seconds. Notably, the DS flies (n=4) displayed a pronounced reduction in speed when transitioning from 10% to 80% RH. This decline was statistically significant in comparison to their speed distribution at both 10% and 80% RH (p<0.05) (Figure 6A). In contrast, the abrupt humidity shifts had no discernible impact on the locomotor speed of the Sated (n=4), DS ir93a (n=3) and sated Ir93a groups (n=3). Speed distributions remained consistent across both humidity levels and exhibited no statistical significance (p > 0.05) (Figure 6B, 6C and 6D). These findings underscore the differential responsiveness of DS flies to rapid humidity changes within the controlled environment.

**Figure 6:**
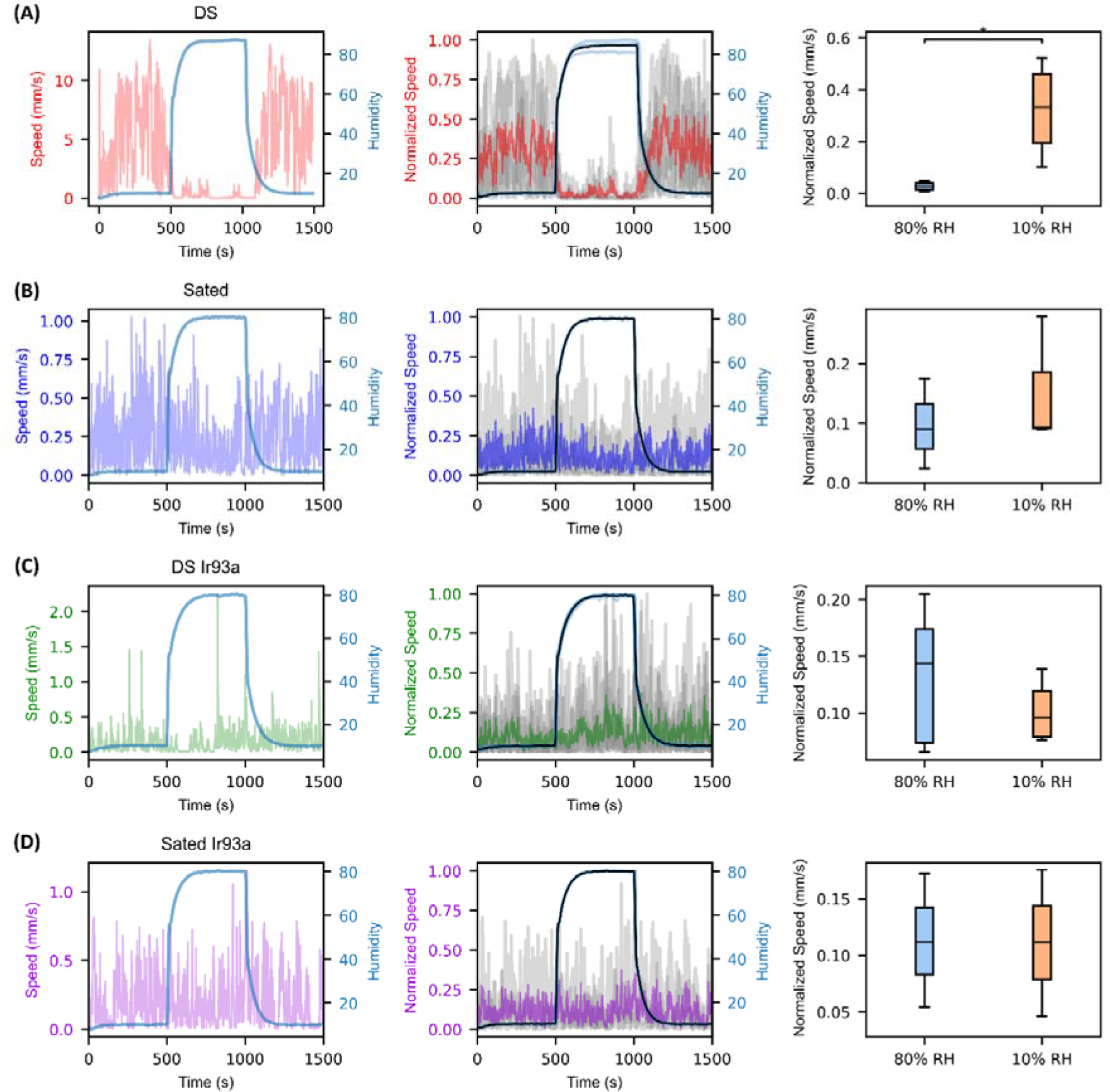
Forced humidity experiments. Speed vs time graph of an individual fly of DS, sated, DS Ir93a and sated Ir93a group respectively for a step humidity function. Each humidity setpoint was maintained for a duration of 600 seconds. Normalized speed vs time graph for all the individual flies of the four groups. Red, blue, green, and violet plot represents the mean normalized speed for DS, sated, DS Ir93a and sated Ir93a group, respectively. The black curve shows the mean humidity for the given group. Distribution of average speed of flies for each trial at 80% and 10% relative humidity (A-D). (* = p<0.05)

## Discussion

This study is the first to elucidate humidity preference of *D. melanogaster* across a continuous 10-80% RH range by developing a novel dynamic humidity arena characterized by precise humidity control and minimal error rates. The experimental results show that desiccated and starved *w*^*1118*^ flies have a preference towards 65-70% RH and exhibit greater speed under conditions of low humidity as compared to high humidity levels, whereas no humidity preference or disparity in speed is evident between low and high humidity conditions in sated and Ir93a flies (DS Ir93a and sated Ir93a).

## A novel experimental setup for humidity-guided behaviours

Humidity is a difficult parameter to control. Previous studies on humidity-guided behaviours in animals have relied on saturated salt solutions that yield stable humidities at fixed values^18^ or ‘dry’ or ‘moist’ air streams with unspecified values of relative humidity^7–13,15,19,20^. While this kind of assays has proven useful in delineating the cells and genes involved in hygrosensation, the limited humidity range tested does not match the full ecological range the fly is exposed, and they cannot be used to precisely quantify the preferred humidity. Therefore, to study the response in all possible ranges of humidity we developed a dynamic humidity arena by integrating the fly-on-a-ball setup with a closed loop humidity delivery system^16^.

The dynamic humidity arena developed in this study can provide a continuous humidity stimulus ranging from 10% to 80% RH. The system comprises two major components: a tracking unit and a humidity delivery unit. The tracking unit continuously monitors the fly’s trajectory, while the humidity delivery unit is responsible for generating and delivering the required humidity levels to the fly. To achieve the desired humidity, the system blends dry and humid air streams in precise proportions using two valves. This humidity delivery system can operate independently, allowing for the maintenance of various humidity set points. Alternatively, it can be integrated with the tracking unit to create a closed-loop system, where humidity levels are adjusted based on the fly’s trajectory. This innovative setup enables the creation of intricate humidity landscapes, facilitating the study of humidity-driven navigation behaviours in *D. melanogaster* as well as other insects.

The observed latency for humidity change was approximately 83 seconds. While more aggressive tuning of PID parameters had the potential to further curtail settling time, it also led to a significant overshoot. To achieve a balance between efficiency and experimental interference, we optimized the flow rate of humidified air delivered to the fly at 1 litre per minute. This configuration not only minimized the interference associated with a direct head-on wind but also enabled consistent humidity control within a range of 10-80% RH.

## State-dependent humidity-preference in *D. melanogaster*

The experimental results demonstrates that DS flies exhibit greater speed under conditions of low humidity as compared to high humidity levels, whereas no disparity in speed is discernible between low and high humidity conditions in sated and Ir93a flies.

The process of desiccation and starvation altered the internal state of both DS and DS Ir93a flies and acted as a catalyst to engage them in extended exploration and cover substantial distances within the arena. However, only DS flies possessed functional hygrosensory receptors, enabling them to precisely locate and favour the 65-70% RH range, where they spent most of their time. In contrast, DS Ir93a flies exhibited a random distribution of humidity preferences, lacking the ability to discern a specific RH level. Sated and sated Ir93a flies, which did not experience desiccation and starvation, exhibited reduced motivation for exploration, this was reflected in their limited arena coverage. Due to limited exploration, these flies remained closer to the initial humidity they were introduced in and created a bimodal distribution for humidity preference depending on the type of humidity map used during the experiment.

Therefore, desiccation and starvation induced alterations in the fly’s internal state that motivated them to seek out a humidity level conducive to their survival, a preference that was most pronounced in DS flies with intact hygrosensory receptors. HRNs express multiple neuropeptide receptors related to state-dependence so it is possible that modulation of the sensitivity already on the primary sensory neurons underly this state-dependent humidity preference^21^.

The results should be viewed under the context that the experiments were performed in a condition where other stimulus such as vision and olfaction were deliberately avoided, to emphasize response solely due to humidity changes, which seldom happens in a natural environment. The observed response of the flies to low humidity, characterized by an increase in speed, suggests an adaptive behaviour aimed at relocating from an unfavourable environment to a more hospitable one. Therefore, speed should not be directly correlated with humidity levels; rather, it should be considered as a responsive behaviour contingent upon the internal state of the fly, or the environment considered favourable by the fly.

## Concluding remarks

In summary, this study describes a novel behaviour arena to study humidity-guided behaviours. We use this arena to provide insights into how humidity affects the behaviour in DS, sated and humidity impaired flies. The unique humidity preference observed in DS flies suggest a nuanced response to environmental humidity conditions. These findings enhance our understanding of the complex relation between humidity and behaviour in this model organism and serves as a stepping stone for further research in the field of hygrosensation.

## ACKNOWLEDGMENTS

The authors thank Kristina Corthals and Marcus Stensmyr for comments on the manuscript and Ola Jakobsson for technical help with valves. This project was funded by the Swedish Research Council, the Crafoord foundation, Ollie and Elof Ericssons stiftelse, Magnus Bergvalls stiftelse, Kungliga fysiografiska sällskapet and the Jeanssons foundation.

## Author Contribution

AE and GG designed the study. GG built the behaviour apparatus, performed all experiments and data analysis. NN assisted in building the behaviour apparatus and data analysis. AE and GG wrote the manuscript with input from NN.

## Data availability

Data and scripts used for analysis can be found on https://github.com/enjinlab/DynamicHumidityArena.

## Materials and methods

### Fly strains

We employed *w*^*1118*^ (Bloomington id: 5905) flies and *Ir93a*^*MI05555*^ (Bloomington id: 42090) mutant flies. These fly strains were maintained at a constant temperature of 25 °C. The humidity levels inside the vials housing the flies varied between 70% and 90% RH, depending upon the proximity to the food source. Upon eclosion, flies were collected and subsequently acclimated to room temperature conditions (21-23 °C) to facilitate habituation. The flies utilized in the experiments exhibited an age range of approximately 7 to 14 days. Prior to the experiments, the flies were starved and desiccated at 5% RH at 21-22 °C for a duration of 4 hours. We also incorporated fully satiated flies into our study, excluding any desiccation or starvation treatment.

### Desiccation and Fly mounting

An insect pin was bent 90 degrees, 1-2 mm from the tip, forming a hook. The pin was then coated with UV curing glue (HelioBond) by collecting a small droplet at the tip. Flies were anesthetized on ice, transferred to a chilled metal plate, and examined under a stereomicroscope with suitable magnification to select healthy specimens. Flies were positioned on their legs with the thorax accessible and attached to the adhesive-coated pin. The adhesive was cured using a handheld UV lamp for 5-6 seconds. After curing, the pin was transferred to a sponge, which covered the opening of the desiccation chamber. The desiccation chamber was fashioned out of a cylindrical container with a tubing inlet for dry air. The internal humidity of the chamber was maintained at 5-6% RH.

### Dynamic humidity arena

Our behavioural apparatus, the dynamic humidity arena, was adapted from the fly-on-the-ball setup^16^. The arena, built on an optical bench, featured a 9 mm ball carved out of plastic (FR-4618, General Plastics) suspended atop a 3D-printed ball holder via air stream. Non-repetitive patterns were drawn on the ball’s surface with a waterproof marker and dried to prevent smudging. An adjustable flow meter was used to regulate the airstream from the ball holder to maintain smooth suspension by eliminating wobbling of the plastic ball. A high-speed camera (Basler acA-1920) with a 150-fps frame rate was used to capture the rotation of the ball. A micromanipulator, mounted on a pedestal post near the ball holder was used to precisely position a tethered fly on the suspended ball.

Once the fly was positioned on top of the ball, the rotation of the ball was then dependent on the walking pattern of the fly. Utilizing images of the rotating ball captured by the camera, we employed FicTrack, an open-source software library^17^, allowing real-time trajectory tracking to reconstruct the fictive path of the walking fly on the patterned sphere.

To tightly control the humidity experienced by the fly, we developed a humidity stimulus delivery system comprising two Bronkhorst LOW-ΔP-FLOW F-201EV flow meters, which regulated the flow of dry and moist air. Mixing these air streams in different proportions, humidity levels from 7-86% RH was achieved in the resultant air stream, though for experiments we capped the RH between 10-80% RH to maintain consistency and reproducibility. The mixed air was delivered to the tethered fly mounted on the ball through a pipette nozzle having a tip diameter of approximately 2 mm. The flow rate was maintained at 1 L/min. A humidity and temperature sensor (Sensirion SHT40 digital humidity sensor) was attached in close proximity to the mounted fly’s antenna which recorded the delivered humidity, serving as feedback for controlling the system. This created a closed-loop humidity delivery system which was further integrated with the fly-on-the-ball setup to form the dynamic humidity arena.

## Experimental Protocol

To investigate humidity preference, flies in the DS and DS Ir93a groups underwent tethering, desiccation, and a four-hour starvation period before each trial. Sated and sated Ir93a group flies were directly tethered from the food source. Each tethered fly then explored the dynamic humidity arena for approximately one hour, with humidity levels determined by their distance from the starting position. Two humidity mapping strategies were employed during the experiment: a low-to-high humidity map with progressive increases over 50 mm, followed by corresponding decreases, and a high-to-low map with the inverse progression. Half of each group experienced the low-to-high map, and the other half the high-to-low map. Throughout the observation, parameters such as humidity, temperature, XY coordinates, and sex were recorded. For forced humidity experiments, DS and mutant flies were tethered, desiccated, and starved for four hours, while sated group flies were taken directly from food. These flies were then introduced to the dynamic humidity arena and exposed to a stepwise humidity function ranging from 10% to 80% RH, with each humidity level maintained for 500 seconds. The entire experiment spanned 1500 seconds.

## Data Analysis

Data analysis for the experimental results was conducted using custom Python programs. Throughout data acquisition, rigorous monitoring of tracking and humidity stimuli was maintained, and any errors in humidity delivery or tracking led to experiment termination. Data from such aborted experiments were excluded from the analysis. X and Y coordinates, along with humidity, temperature, and time, were extracted from the saved data files generated during the experiments.

An exponential weighted moving average filter with a span of 50 was applied to the data to remove noise and facilitate further processing. Using the positional coordinates (X and Y), the Euclidean distance between each data point was calculated. Cumulative distance was obtained by the addition of distance values over time. Speed between each data point was then computed using the acquired distance and time interval values recorded during the experiments.

Humidity preference among DS, sated, DS *r93a and sated* ir93a flies, were analysed by constructing histograms presenting the combined humidity values from each group, categorized by the employed humidity map. To delve into individual preferences, we extracted humidity peaks from each trial’s histogram and represented them as boxplots. Statistical comparisons of humidity preferences across the three groups were conducted using the Mann-Whitney U test, with Bonferroni correction for multiple comparisons.

Speed data was grouped into 10% RH intervals, and the mean speed was computed for each interval. These results were visualized using scatter plots to examine the relationship between humidity and speed. To account for potential confounding factors such as desiccation state, sex, and group (DS, sated, DS Ir93a *or sated* Ir93a), we employed a mixed-effects model. To prepare the data for the model, a custom Python function was used to merge the data files for the three groups, with sex determined based on the initial character in the file names (m for male and f for female). Box-Cox transformation to the speed data was applied to improve normality. Using the Statsmodels library, a mixed-effects model was built that regressed speed against humidity, group, sex, and their interactions. The identifier column which labelled individual trial served as a grouping variable. Model summaries and coefficients were stored for interpretation. Speed profiles were generated by specifying humidity values and obtained model coefficients for different group-sex combinations.

In the forced humidity experiments, fly speeds were categorized into two humidity levels: 10% and 80% RH for each trial within the group. Speed data were normalized to account for trial-to-trial variations within the group. Mean speeds at these two humidity levels were then calculated and depicted using box plots. Given the non-normal distribution of the data, we employed a non-parametric Mann-Whitney U test followed by Bonferroni correction to assess whether there was a statistically significant difference in fly speed between the 10% and 80% RH conditions within the given group.

