## Supplemental figure for "A dynamic humidity arena to explore humidity related behaviours in insects"

**Supplementary figure**

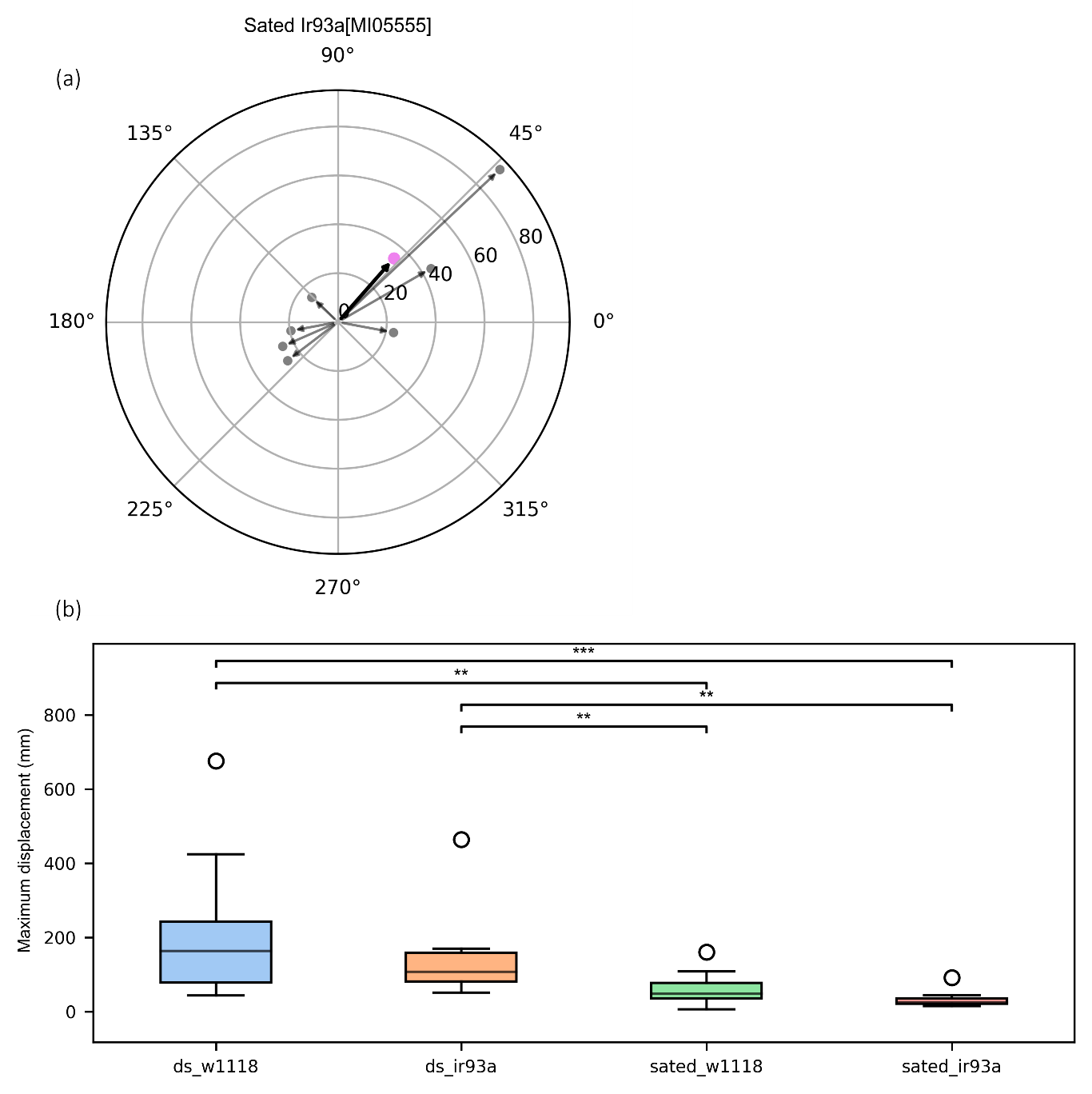

***Supplementary figure 1:*** *Distribution of maximum displacement in DS, DS Ir93a, sated and sated Ir93a group. The significant difference in maximum displacement between the desiccated and starved flies and sated flies suggest that the act of desiccation and starvation caused a change in the internal state of the fly compelling it to explore further in search of a favourable environment* *(*** = p<0.001, ** = p<0.01, * = p<0.05)*
